# A Solution Method for Predictive Simulations in a Stochastic Environment

**DOI:** 10.1101/793588

**Authors:** Anne D. Koelewijn, Antonie J. van den Bogert

## Abstract

This short communication describes a method to solve predictive simulations of human movements in a stochastic environment using a collocation method. The optimization is performed over multiple noisy episodes of the trajectory, instead of a single episode in a deterministic environment. Each episode used the same control parameters. The method was verified on a torque-driven pendulum swing-up problem. A different optimal trajectory was found in a stochastic environment than in the deterministic environment. Secondly, it was applied to gait to show its application in human movements. We show that nonzero minimum foot clearance during swing is energetically optimal in a stochastic environment. The amount of clearance increased with the noise amplitude.

## 1. Introduction

Predictive gait simulations have been used to show the effect of interventions on gait (Koelewijn and van den Bogert, 2016; Dembia et al., 2017; Van den Bogert et al., 2012; Dorn et al., 2015; Miller et al., 2013). However, they fail to predict several key patterns of human gait (Ackermann and van den Bogert, 2010). For example, predictions of gait of persons with a transtibial amputation (TTA gait) failed to predict co-contraction in the thigh on the prosthesis side (Koelewijn and van den Bogert, 2016), while this is observed in electromyography studies of TTA gait (Isakov et al., 2000; Powers et al., 1998). This seemingly sub-optimal behavior increases limb stiffness and may improve stability against perturbations (Powers et al., 1998).

We aim to include uncertainty into predictive simulations of human movements, since several studies have highlighted the importance of environmental uncertainty when choosing movement patterns (Hiley and Yeadon, 2013; Kim and Collins, 2015; Donelan et al., 2004), but uncertainty is currently ignored in simulations. When it is not, the stochastic differential equations, or the stochastic Hamilton-Jacobi-Bellmann equation, should be solved. This is difficult and the problem becomes intractable for nonlinear, high dimensional systems (Kappen, 2005), such as musculoskeletal models. Also, the certainty equivalence principle does not hold for nonlinear systems Todorov (2005).

Instead, the stochastic environment could be estimated. Monte Carlo methods are used to estimate the solution of trajectory optimization problems with uncertainty (Tiesler et al., 2012; Sandu et al., 2006; Matsuno et al., 2015). A faster approach is to use generalized polynomial chaos, where the uncertainty is approximated with a combination of stochastic basis functions. This problem can be solved with the stochastic Galerkin method or stochastic collocation (Tiesler et al., 2012; Sandu et al., 2006).

In this paper, a method for solving trajectory optimization problems in a stochastic environment is applied to human movements. The method can be seen as generalized polynomial chaos with basis functions of the first order, solved using stochastic collocation. The approach will be verified on a classic one degree-of-freedom pendulum swing-up problem. Then, it will be applied to a simple gait problem, specifically minimum foot clearance (MFC), which is the minimum foot-ground clearance during swing. It is equal to about 1.29 cm in normal gait (Winter, 1992), while zero MFC would likely be most energy efficient, especially in a deterministic environment. We aim to show that nonzero minimum foot clearance (MFC) is optimal in a stochastic environment.

## 2. General Problem Description and Solution Method

We aim to solve trajectory optimization problems in a stochastic environment, meaning that the dynamics, with state *x* and input *u*, are dependent on some noise, *ε*:

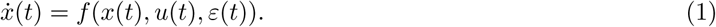

This noise causes unstable systems, such as human models, to deviate from the desired path, requiring closed-loop control. Additionally, the objective cannot be determined exactly from one episode, since its value will depend on the noise sample.

In theory, an infinite number of episodes is required to accurately estimate the objective, which is computationally intractable. Therefore, similar to a Monte Carlo approach, an estimation will be made with a finite, sufficiently large, number of episodes of the trajectory. Each episode uses the same open-loop inputs and the same closed-loop control parameters. The objective is estimated using the average over the episodes.

Instead of using forward simulations of the complete trajectory, the trajectory optimization problem is solved using a collocation method, a fast method that can use the analytic gradient of the dynamics (Van den Bogert et al., 2011). The optimization is performed over the states of the *N* collocation points of all *N*_*s*_ episodes, and the control parameters: *X* = [*x*_1_(1) … *x*_1_(*N*) … *x*_*N*_*s*__ (1) … *x*_*N*_*s*__ (*N*), *u*_0_(1) … *u*_0_(*N*), *K*]^*T*^, where *x*_*j*_(*n*) denotes the state at collocation point *n* in episode *j*, *u*_0_(*n*) the open-loop input at collocation point *n*, and *K* represents the closed-loop control parameters. This yields the following problem description:

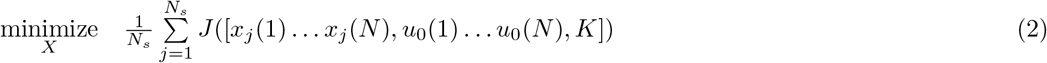

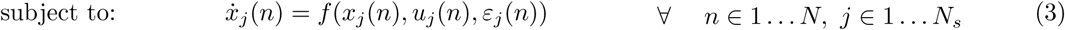

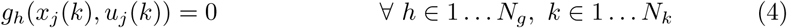

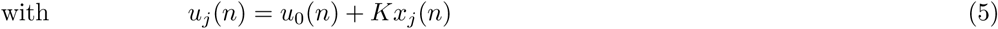

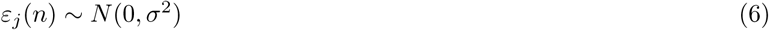

where *g*_*h*_ describes task constraint *h*. The estimated objective is described in equation 2. Equation 3 represents the dynamics constraints, which should be met for all collocation points and all episodes. Noise will be added at each collocation point. Equation 4 represents the task constraints, which are applied at *N*_*k*_ collocation points *k*. Equation 5 denotes the feedback law, with a proportional and a derivative feedback gain. Note that *u*_0_(*t*) is the open loop control term which is identical for all episodes.

This approach will be applied to two problems. The first problem is a classical pendulum swing-up, where the goal is to show that this method is able to find a different optimal trajectory in a stochastic environment. The second problem highlights the applicability to gait by looking at the MFC in a deterministic and stochastic environment.

## 3. Swing-Up Problem

In the swing-up problem, the task is to move an arm from the downward position to the upward position while minimizing an objective related to energy (see figure 1). Since the upward position is an unstable equilibrium, the optimal trajectory will swing-up faster and using more active control due to the higher cost (Todorov, 2011).

**Figure 1:**
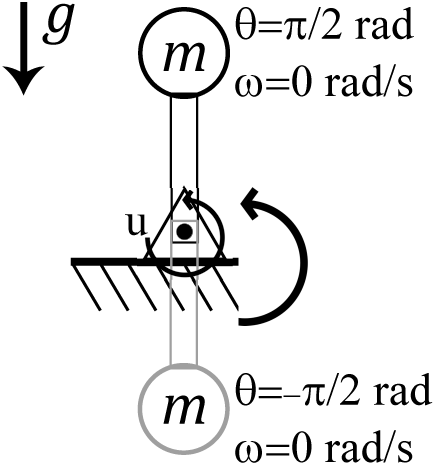
Arm used for the swing-up problem. The goal was to move this arm from the downward to the upward position in ten seconds, while minimizing the squared torque.

### 3.1. Methods

Figure 1 describes the problem that is solved. The arm dynamics, with angle *θ* and angular velocity *ω*, were as follows:

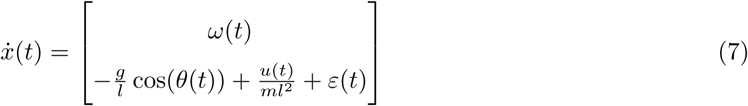

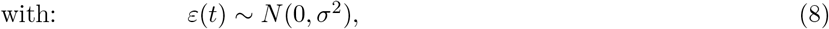

where *g* = 9.81 m/s denotes gravity, *m* = 2 kg the mass of the pendulum, and *l* = 0.6 m the location of the center of mass. The input to the system was the torque *u* at the base of the arm. In the stochastic environment, normally distributed noise was added to the angular acceleration with a standard deviation *σ*. Closed-loop control, *u*(*t*), was determined using the angle and angular velocity, according to equation 5.

The objective was to minimize the square torque, which is related to energy, yielding the following trajectory optimization problem:

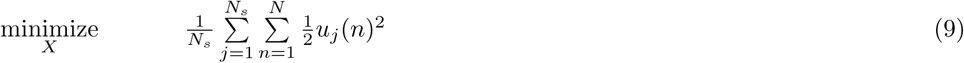

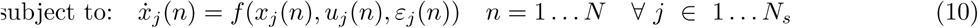

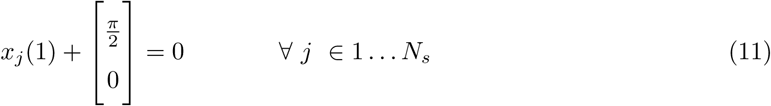

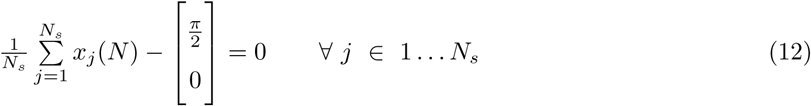

where the task constraint on the final position (equation 12) was averaged over all episodes, since the final position could not be controlled exactly in the stochastic environment. The supplementary material describes the pipeline to find the solutions. All code is available here (Koelewijn, 2019).

### 3.2. Results and Discussion

Figure 2 (upper) shows the mean optimal trajectories for different noise amplitudes, averaged over 10 solutions to account for variation due to noise. As expected, a different trajectory was optimal in the stochastic environment than in the deterministic environment. The timing of the swing up changed such that the final swing-up (between 8 and 10 seconds) occurred later for a larger noise amplitude.

**Figure 2:**
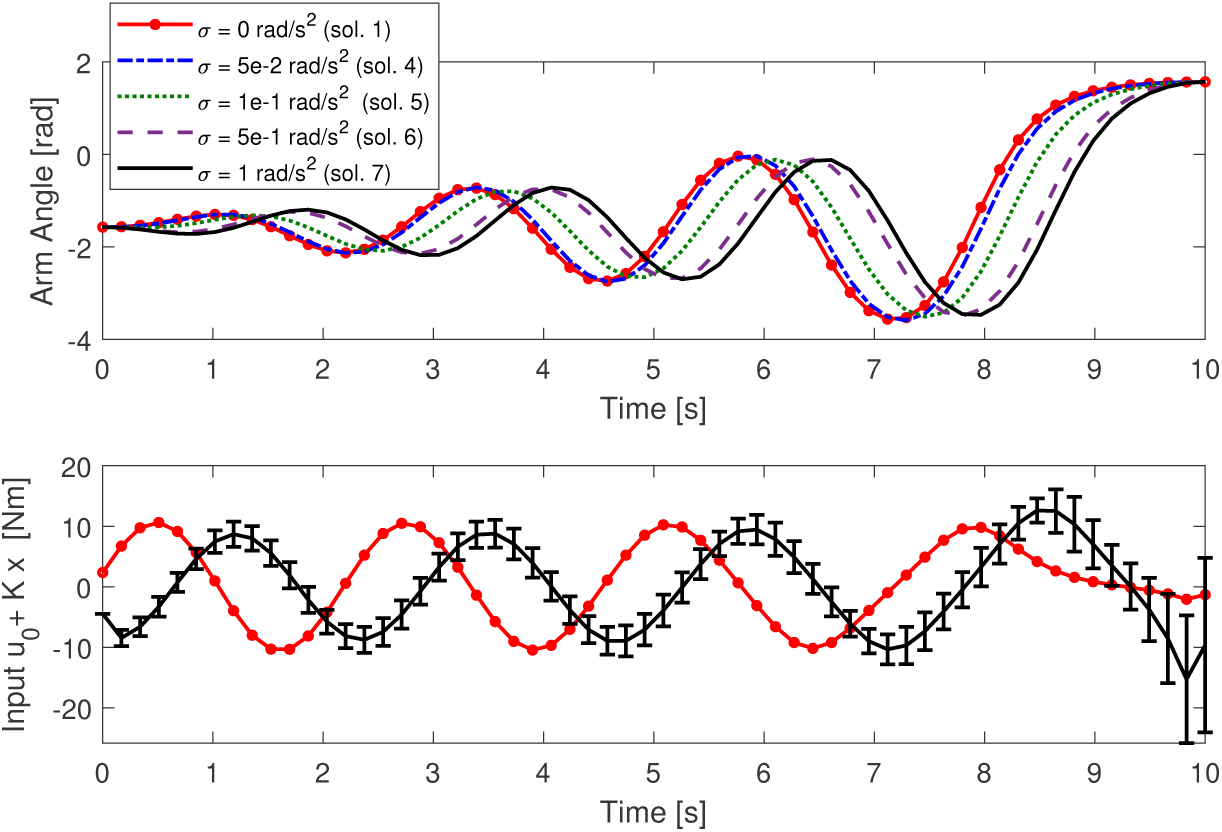
Optimal swing-up trajectories, mean of all episodes (upper graph), and control torque, mean and standard deviation, (lower graph) for solutions 1, 4 - 7 (denoted as sol.) according to the supplementary material, table 1.

Compared to the deterministic environment, the timing of the control input for *σ* = 1 rad/s^2^ changed similarly (figure 2, lower). Additionally, the final swing-up in the stochastic environment required more control than in the deterministic environment, and had the largest variation in control as well (vertical bars). This indicates a more active control approach in the stochastic environment compared to a passive approach in the deterministic environment, where gravity is used to brake the pendulum. The supplementary material describes additional results of the pendulum swing-up to show that with seven episodes the episode is already estimated correctly, and that closed-loop control reduced the variability of the result, while the conclusion was the same without closed-loop control.

## 4. Minimum Foot Clearance in Gait

The second problem aims to highlight the application of the solution method to human movement, specifically gait. We aim to investigate if a nonzero MFC would be energetically optimal in a stochastic environment.

### 4.1. Methods

A torque-controlled, sagittal plane model with nine degrees of freedom was used (see figure 3). The trunk position and orientation were unactuated, while the hip, knee and ankle angles were torque-controlled, with an open loop and a closed loop part that used the position and velocity of the same degree of freedom (see equation 5).

**Figure 3:**
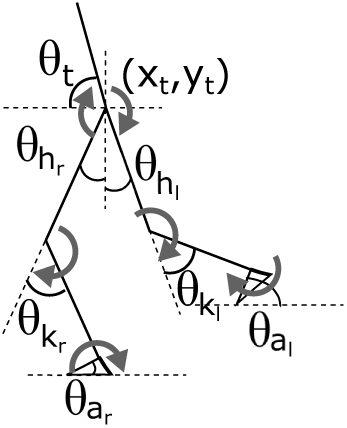
Overview of the sagittal plane human model. All nine degrees of freedom are shown, as well as the locations of the torque inputs.

The objective was to minimize square torque and to minimize a tracking error between the simulation and 11 normal walking variables (Winter, 1991): three joint angles, the ground reaction forces (GRFs) and the gait cycle duration:

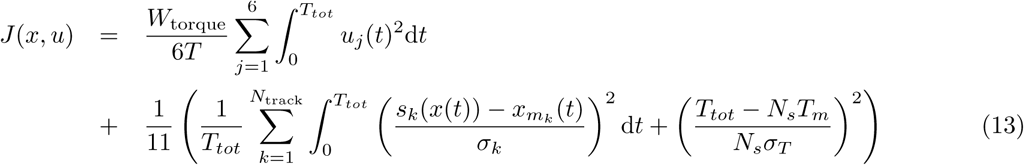

where *s*_*k*_(*x*(*t*)) denoted the normal joint angles and GRFs, with standard deviation *σ*_*k*_, *T*_*tot*_ the total duration of the movement, *T*_*m*_ the tracked duration with standard deviation *σ*_*T*_, and *N*_*s*_ the number of episodes (gait cycles), *N*_*s*_. *W*_torque_ = 0.001 was used to create a realistic gait cycle. A similar objective, minimizing square muscle activation, was shown to predict human gait well (Van den Bogert et al., 2012; Koelewijn and van den Bogert, 2016).

Each episode consisted of a half gait cycle, assuming left-right symmetry. The task constraint was a translation of the horizontal trunk position, *x*_*t*_, for a fixed speed and a periodicity constraint between the first collocation point of the first episode and the last collocation point of the last, tenth, episode, yielding the following task constraints:

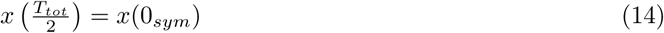

and

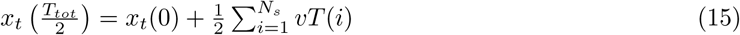

where *T*(*i*) denoted the duration of gait cycle *i*, the subscript *sym* denoted that the joint angles and angular velocities were switched between the legs, and *v* is the speed. Dynamics were constrained between the other episodes to simulate a series of steps. 10 gait cycles were chosen to limit computational power. The supplementary material details the solution pipeline.

### 4.2. Results and Discussion

Figure 4 shows the average clearance of the heel and toe for noise amplitudes from 0 Nm to 100 Nm. During the stance phase, the clearance is negative, due to the ground penetrating contact model that was used (see Koelewijn and van den Bogert (2016)). The graphs on the right show the part of the swing phase where the MFC occurs. In this phase, the MFC was exactly zero with deterministic dynamic and increased with the noise amplitude.

**Figure 4:**
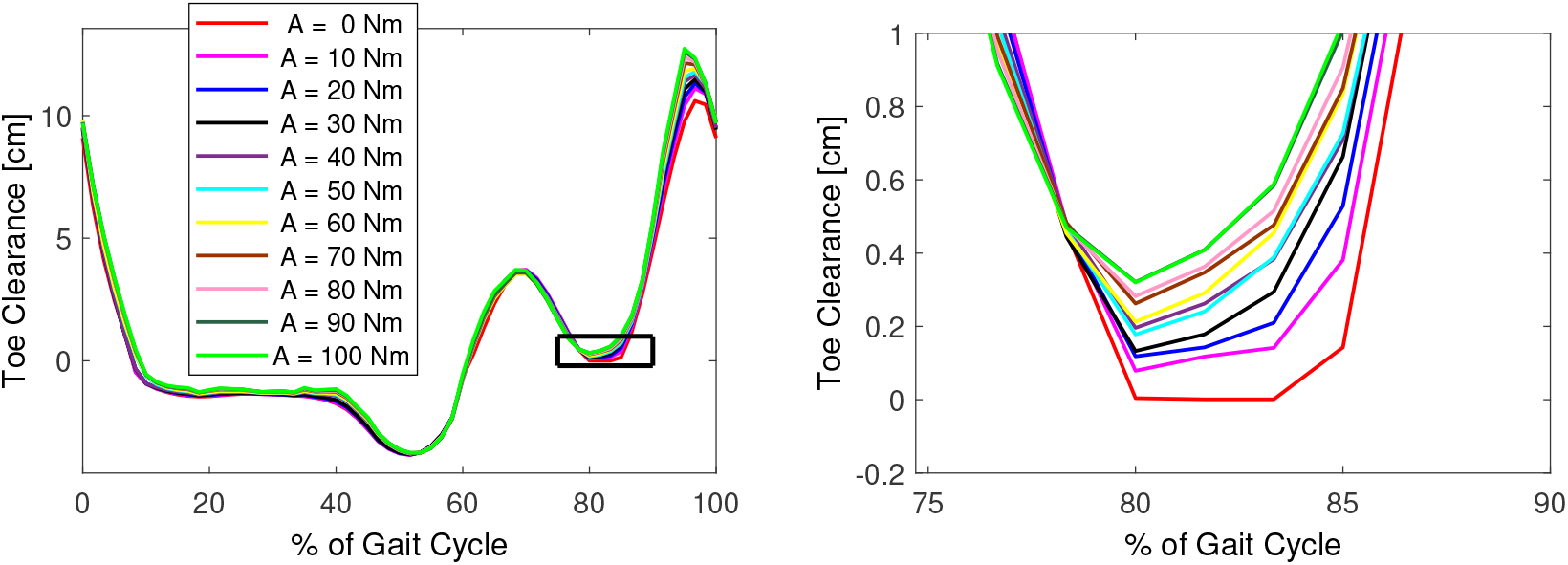
Average heel (top figures) and toe clearance over 10 gait cycles for solutions with different noise amplitudes. One can see that with an increasing noise amplitude, the clearance of the heel and toe increases.

These results show that in a stochastic environment, nonzero MFC is energetically optimal and that the MFC increased with the amplitude of the noise. Inspection of all gait cycles showed that the MFC was exactly zero at one instance, meaning that the average ground clearance was just high enough to prevent scuffing, similar to an observation in an experimental study (Wu and Kuo, 2016). The MFC was still lower than in normal gait Winter (1992). This is likely due to the noise model, which is too different from noise in human gait due to neural control or uneven ground to yield a realistic answer.

The closed-loop controller in this problem was a proportional-derivative controller with the same two gains in each joint. This was done to limit the problem’s search space. A more complex controller will likely yield a lower objective, but requires a larger number of episodes. Ten episodes were solved in approximately four hours on a standard laptop.

## 5. Conclusion

This paper solved trajectory optimization problems of human movements in a stochastic environment. These simulations can potentially explain human motor control behavior which cannot be explained in a deterministic environment. We verified our solution method on a pendulum swing-up problem. To show the applicability to human movement, we showed that nonzero MFC was optimal in gait in a stochastic environment.

### 5.1. Future Work

The motivation of this work was to be able to account for uncertainty in trajectory optimization problems to improve predictive simulations of gait. Specifically, we aim to understand co-contraction of muscles in TTA gait. Co-contraction is often described as inefficient, because it does not produce any useful work while it requires energy (Hogan, 1985). However, we hypothesize that co-contraction is optimal in an environment with uncertainty.

To test this hypothesis, we aim to apply the approach that was introduced in this paper. When it is combined with a musculoskeletal simulation, we can see if co-contraction (a feedforward approach) is more energy efficient for certain movement tasks than a reactive approach. One of the applications will be TTA gait. Care should be taken to create a realistic model of uncertainty in human movements. Additionally, due to the large number of states and optimization variables in the musculoskeletal model, this problem could be difficult to solve.

## Acknowledgements

This research was supported by the National Science Foundation under Grant No. 1344954 and by a Graduate Scholarship from the Parker-Hannifin Corporation.

