## Supplementary Material for "A Solution Method for Predictive Simulations in a Stochastic Environment"

#### Explanation of terms and variables

This section explains the terms and variables that are used in this work. The variables are listed in table 1 and the terms in table 2.

Table 1: List of variables

| | |
|---|---|
| $x$ | Dynamics state |
| $u$ | Control input |
| $\varepsilon$ | Noise in system, environmental or internal |
| $N$ | Number of collocation points |
| $N_e$ | Number of episodes |
| $n$ | Collocation point |
| $j$ | Episode |
| $u_0$ | Time dependent, open loop control input |
| $K$ | Closed-loop control parameters |
| $h$ | Task constraint |
| $N_g$ | Number of task constraints |
| $N_k$ | Number of time points where constraint is applied |
| $k$ | Time point where constraint is applied |
| $\sigma$ | Standard deviation of Gaussian noise |
| $A$ | Amplitude of uniform noise |

#### Supplementary Methods - Pendulum Swing-up

Table 3 describes the trajectory optimization problems that were solved. Direct collocation was used with a midpoint Euler approximation for the dynamics [1]. $N = 60$ collocation points, so a time step of 0.16 s, were used and $N_s = 30$ episodes. The deterministic problem was solved from an initial guess at the middle of the search space. The problem was repeated with 100 random initial guesses. The objective of the solution from

Table 2: List of terms

| | |
|---|---|
| Objective | The function that is minimized |
| Noise sample | One instance of the noise sampled from the mean and amplitude/standard deviation |
| Episode | One instance of the motion with a specific noise sample |
| Collocation point | Time instance where the dynamics constraints are applied |
| Problem | Complete optimization problem with all episodes |
| Solution | Set of states and inputs that optimizes the objective |

Table 3: Description of the problems that were solved and their initial guess.

| Number | Environment | Standard deviation [rad/$s^2$] | Initial Guess |
|---|---|---|---|
| 1 | Deterministic | - | Middle of search space |
| 2 | Stochastic | 0.001 | Solution 1 |
| 3 | Stochastic | 0.01 | Solution 2 |
| 4 | Stochastic | 0.05 | Solution 3 |
| 5 | Stochastic | 0.1 | Solution 4 |
| 6 | Stochastic | 0.5 | Solution 5 |
| 7 | Stochastic | 1 | Solution 6 |

the middle initial guess was the lowest that was found of these random guesses. The deterministic solution was then used as initial guess for the stochastic problem with the lowest standard deviation. This process was repeated for increasingly large standard deviations. Noise was sampled once for the series of problems and weighted according to the standard deviation of the current problem. The deterministic solution was compared against the stochastic solutions with a standard deviation of 0.05 rad/s$^2$ or larger.

$3N$ variables were optimized in the deterministic environment, two states and one control input at each collocation point, with $2N$ equality constraints. In the stochastic environment, there were $2NN_s + N + 2$ optimization variables, for optimization variables $X = [x_1(1) \dots x_1(N),\ x_2(1) \dots x_{N_s}(N),\ u_0(1) \dots u_0(N),\ K_P,\ K_D]^T$. The number of constraints was equal to $2NN_s$.

The objective function and the dynamics and task constraints, and their derivatives with respect to $x$, $\dot{x}$ and $u$ were coded in MATLAB (Mathworks, Natick, MA, USA). All optimization problems were solved using IPOPT 3.11.0 [2].

### Supplementary Results - Pendulum Swing-Up

#### Convergence Analysis

Figure 1 was used for the convergence analysis. It shows the objective of the optimal solution as a function of the number of episodes. When more than 20 trajectories were used, the objective converged to a value of 26.5 (Nm)$^2$ and did not change more than 1% compared to the objective found with 100 episodes, the largest number of episodes used. Also, the variation of the objective remained lower than 0.2 Nm using 20 or more episodes.

The value of the objective function did not change more than 1% when at least 20 episodes were used. Also, the standard deviation over 10 solutions remained 0.2 Nm. Figure 1 shows that already with seven samples, a good estimate of the optimal objective in the stochastic environment was found. With five or more episodes, the trend is visible that was shown in figure 2 of the manuscript, that the swing-up occured

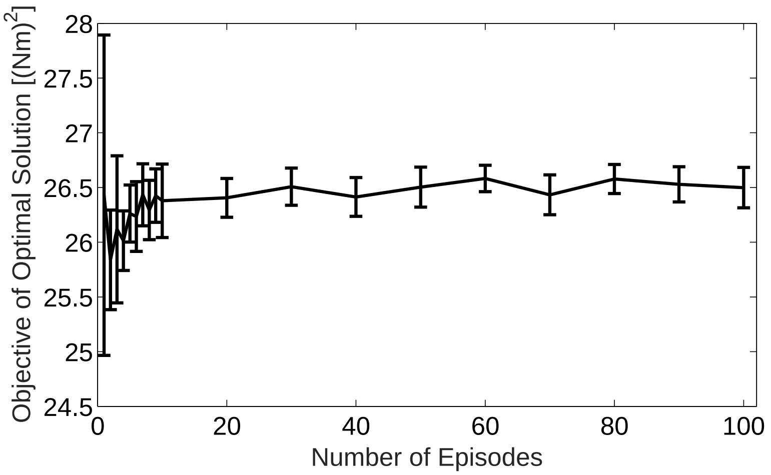


Figure 1: Convergence analysis of the objective of the optimal solution. Convergence occurred after 20 samples to a value of 26.5 $(\mathrm{Nm})^2$.

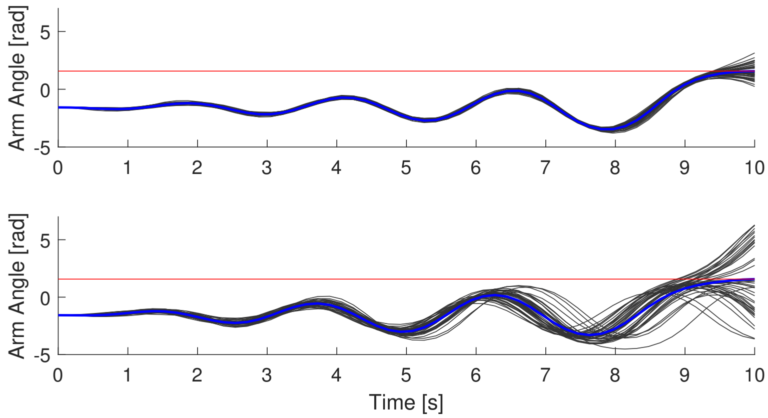


Figure 2: Comparison of the trajectories found with feedback control (top figure) and without feedback control (bottom figure) for a standard deviation of 1 rad/s$^2$. The black line shows the average trajectory, while the (40) black lines show each individual trajectory and the red line shows the 90 degrees.

increasingly late with a larger standard deviation.

#### Closed-loop Control

Figure 2 shows the trajectories that were found using open-loop and closed-loop control (upper) and with open-loop control only. One can see that the variation was much higher using only open-loop control, such that certain episodes did not swing-up, while others overshot and ended up in the downward position.

The combination of closed-loop and open-loop control decreased the variability of the results, as can be seen in figure 2. However, the conclusion was the same, since the average trajectory without feedback control also swung up later in the stochastic environment than in the deterministic environment. The stability of the system with these feedback gains in the upright position was checked by linearizing the system around the upright position and checking if the real part of the eigenvalues were negative. Stable feedback gains were found for all problems with more than 10 episodes for all standard deviations, while solutions with fewer episodes were stable or marginally stable. This suggests that this method could also be used in robotics, to simultaneously find an optimal trajectory and a controller in one optimization procedure. With a conventional approach in a deterministic environment, only an optimal trajectory is found. The controller is found separately, for example by linearizing around the trajectory.

### Solution Method - Gait Study

The optimal control problem was solved for 10 episodes, with 30 collocation points each and a backward Euler formulation. Van den Bogert et al. [3] derived the dynamics equations for the multibody system with the ground contact model (see [4]) using Autolev (Online Dynamics, Sunnyvale, CA). This function was coded in C and compiled as a MEX-function for MATLAB (Mathworks, Natick, MA). Details of the solution method can be found in [5]. The resulting constrained optimization problem was solved using IPOPT 3.11.0 [2].

First, one gait cycle was found with deterministic dynamics ($\varepsilon(t) = 0$). This gait cycle was used as initial guess in the stochastic environment, which was modeled using uniform noise, $\varepsilon(t) \sim U(-A, A)$ at each collocation point (sampling rate around 0.037 s). Uniform noise was used instead of Gaussian noise to avoid outliers, which caused problems during the optimization.

Next, a predictive simulation was found in a stochastic environment with noise amplitude, $A = 0.5$ Nm, followed by 1 Nm, 5 Nm, 10 Nm, and up to 100 Nm with increments of 10 Nm. 7000 optimization variables were optimized in the stochastic environment.

### Supplementary Results - Gait Study

#### Joint Moments

Figure 3 shows the joint moments over 10 gait cycles for the solution with noise amplitude 100 Nm. This figure shows the full gait cycle, while only half was simulated. The second half is added for completion. The values of the torques are much higher than in humans. However, the shape is as expected.

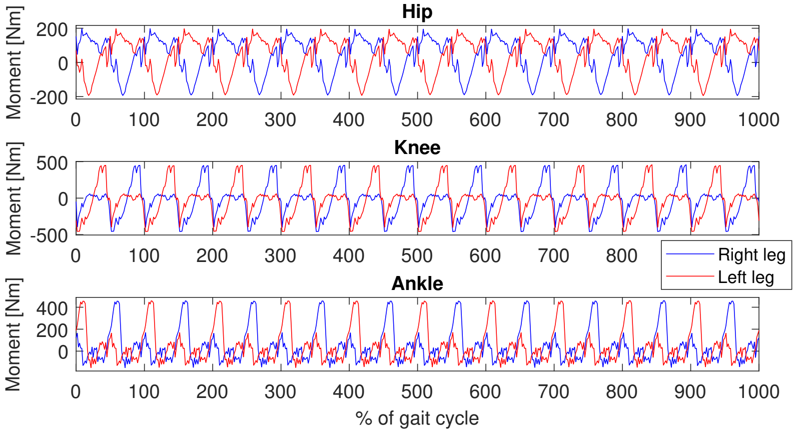


Figure 3: Joint moments in the hip, knee, and ankle joint for the full 10 gait cycles. The horizontal axis gives % of the gait cycle, where each gait cycle has 100%. Winter's convention is used, where extension and plantarflexion are positive.

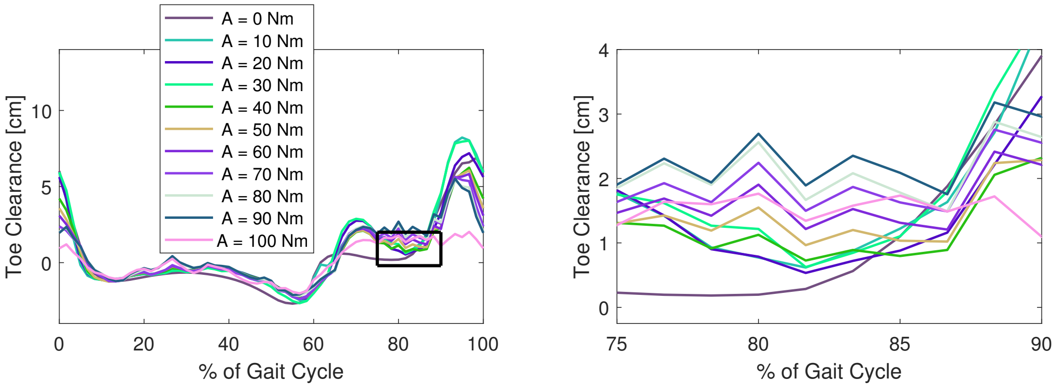


Figure 4: Toe clearance with weight $W_{torque} = 0.01$ for increasing noise amplitudes. Similar to the results presented in the manuscript with a smaller weight on the torque, the foot clearance increases with the size of the noise.

#### Effect of Tracking Weight

The effect of the tracking weight on the solution was analyzed by repeating the solution process but with the weight increased by a factor of 10 to decrease the effect of tracking. Figure 4 shows the toe clearance for this objective. The resulting simulation is much less smooth, but still the same behavior is visible, that the foot clearance increases with noise amplitude.

[5] AD Koelewijn. *Predictive Simulations of Gait and Their Application in Prosthesis Design.* PhD thesis, Cleveland State University, 2018.
